# Introgression of Austrian fire-bellied toads (*Bombina bombina*) into northern German populations confirmed by complete mitochondrial genomes and transcriptome-wide Single Nucleotide Polymorphisms (SNPs)

**DOI:** 10.1101/651695

**Authors:** Binia De Cahsan, Michael V Westbury, Sofia Paraskevopoulou, Hauke Drews, Moritz Ott, Günter Gollmann, Ralph Tiedemann

**Author notes:** Email address: Michael V Westbury –, Sofia Paraskevopoulou - paras, Hauke Drews -, Moritz Ott -, Günter Gollmann. to whom correspondence should be addressed; Binia De Cahsan –, Ralph Tiedemann –.

## Abstract

The fire-bellied toad (*Bombina bombina*) is a small amphibian species inhabiting the lowlands of Central and Eastern Europe. Continual population fragmentation has caused a decrease in genetic diversity and gene flow in margin populations in Northern Germany. Previous research using mitochondrial control region data uncovered a translocation of allochthonous toads from Austria into a single northern German population, Högsdorf. Microsatellite and MHC data also confirmed this introgression, but were limited in revealing the true extent of this translocation for all investigated populations. Here, we utilize complete mitochondrial genomes and nuclear data, in the form of transcriptomes, to unravel the full extent of admixture from this translocation. The increased number of loci enabled us to uncover signs of introgression in four out of the five northern populations investigated. Moreover, as hybrids between *B. bombina* and its sister species, *B. variegata*, have been reported in Austria, we also investigated whether *B. variegata* alleles were translocated into the northern gene pool. We found evidence for hybridization between Austrian *B. bombina* and *B. variegata*, as well as traces of *B. variegata* alleles in one of the five German populations included in this study. These findings present the first reported case of introgressed relict *B. variegata* alleles from the Southern lineage being detected in Northern Germany.

## Introduction

Genetic admixture between different populations is a widespread phenomenon that instantly increases the genetic diversity of a population’s gene pool and can help to promote adaptation to changes in environmental conditions (Michael and Kunte 2017; Janes and Hamilton 2017). The fire-bellied toad (*Bombina bombina*) is a small amphibian species found in the Central and Eastern European lowlands. It mostly inhabits permanent, fish-free freshwater bodies, like shallow stagnant lakes, ponds and swamps (Drews et al. 2011). After the Pleistocene, this species most likely colonized Europe in two distinct expansions out of refugia located northwest of the Black Sea (Hofman et al. 2007; Fijarczyk et al. 2011). The Carpathian Mountains and the Central European middle mountains at the border between Germany and the Czech Republic formed natural barriers that led to a north-south division of the species range. As a consequence, two genetically distinct evolutionary lineages of *B. bombina* emerged (Hofman et al. 2007).

Geographically close populations of *B. bombina* are often genetically distinct. The high site fidelity and limited dispersal capacity of *B. bombina* were proposed as a putative explanation for this phenomenon (Engel 1996; Dolgener et al. 2012; Schröder et al. 2012). This is particularly true for locally restricted and highly fragmented populations at the north-western margin of the species range, which runs from the south of the Swedish province Scania to the Baltic Sea through Denmark and Germany (Schleswig-Holstein, Niedersachsen, Sachsen-Anhalt) (Schröder et al. 2012). In comparison to their counterparts in the central distribution range, margin populations of *B. bombina* are strongly threatened by habitat alterations, pollution and intensive agricultural use (Drews et al. 2011). Continuous human mediated habitat fragmentation has led to a reduction in gene flow and locally isolated populations are genetically depauperated (Schröder et al. 2012). *Bombina bombina* population sizes in Germany have been declining since the middle of the last century, which led to the species conservation status being listed as “critically endangered” within Germany (Kühnel et al. 2009).

Genetic analyses from toads collected in northern Germany in 2006 uncovered a previously undetected introgression of southern lineage (putatively Austrian) genotypes into northern lineage populations. This was likely the result of occasional (and illegal) translocations of toads of southern origin into northern populations (Schröder et al. 2012). In addition, the fire-bellied toad is also known to hybridize with its sister species the yellow-bellied toad (*B. variegata)(Szymura* and Barton 1986). The two species have parapatric distributions due to different habitat preferences. While *B. bombina* prefers larger permanent ponds in lowland areas, *B. variegata* occupies smaller, temporal water bodies found at higher altitudes and warmer temperatures (MacCallum et al. 1998). Despite this, hybridization did occur and narrow hybrid zones are formed where the two species meet (e.g., in Austria, Hungary, and Poland) (Szymura and Barton 1986). So far, traces of a past genetic introgression between these two species have not been detected in northern Germany.

Here, we present both complete mitochondrial genomes and nuclear data, in the form of transcriptomes. Using our data, we searched for signs of introgression between *B. bombina* from Austria and conspecific northern German *B. bombina* populations. We investigated whether mitochondrial introgression from Austria can still be found in the northern German populations or whether it has since disappeared due to random genetic drift or natural selection. Additionally, we investigated whether introgression can also be found in the nuclear genome. Finally, as hybrids between *B. bombina* and *B. variegata* have been reported in Austria, we also investigated whether *B. variegata* alleles were translocated into the northern gene pool by investigating signs of admixture between the yellow-bellied toad and northern *B. bombina* populations.

## Materials and Methods

### Samples

30 wild tadpoles were collected, five from Austria (Lobau, Vienna) in 2016 under permit MA22 - 984143-2015-5 issued by the Municipality of Vienna and 25 from Germany (five each from, Eutin/Röbel, Dannau, Högsdorf, Fehmarn, Neu-Testorf) in 2017 under permits from Landesumweltamt Schleswig-Holstein (See Supplementary Material, Fig. S1). German tadpoles were frozen in liquid nitrogen and stored at −80 °C. The five Austrian tadpoles were transferred into a RNAlater storage solution.

### RNA extraction, mRNA enrichment and cDNA library preparation

We extracted RNA from the tail muscle of the fins, using a modified Trizol isolation protocol in combination with a column based commercial kit (QIAGEN RNeasy Mini Kit) based on the manufacturer’s protocol. Only skin and muscle tissue of the tadpole’s fins were used in order to prevent contamination from gut bacteria during sequencing as much as possible. To further increase RNA yield, we transferred the frozen tadpole tails into a 1 ml Trizol solution and performed a homogenization with a Tissue Lyzer (6 min, 50 HZ). We implemented the column based clean up procedure from the QIAGEN Rneasy Mini Kit as described in their manual. We assessed the quality and quantity of the extracted total RNA on the Agilent Bioanalyzer 2100. We only selected samples with a high RIN number (>8) as well as a high RNA concentration (c > 50 ng/μl) for library preparation. We then performed an mRNA enrichment on these samples with Poly(A) beads using the Bio Scientific NEXTflex Kit following the manufacturer’s protocol. For assessing the quantity of successfully enriched mRNA, we ran a Qubit High Sensitivity RNA Assay. We built cDNA libraries from the enriched mRNA samples with the NEXTflex Rapid Directional RNA-Seq Kit (Bio Scientific) following the manufacturer’s protocol. During the strand specific library preparation, we single indexed each sample and ran a PCR amplification with 15 cycles and an annealing temperature of 65 °C. We performed a final quantity check on the built libraries with a dsDNA High Sensitivity Assay on the Qubit and a quality check on the Bioanalyzer. The cDNA library concentrations ranged from 2.94 ng/μl to 17.4 ng/μl, all with high RIN numbers. We sent the libraries to be sequenced at Novogene in Hong Kong, China, where samples were additionally assessed for sufficient quantity and quality and PE 150 bp shotgun sequenced on three lanes on an Illumina HiSeq.

### Quality filtering and mapping

We trimmed Illumina adapter sequences and poly A tails, only keeping reads with a quality score of at least 25 and a minimum length of 30 bp for all 30 samples using the software Skewer v0.2.2 (Jiang et al. 2014). Overlapping forward and reverse reads we then merged with FLASH v1.2.10 (Magoč and Salzberg 2011) using default parameters. The successfully merged reads and the remaining unmerged reads were mapped separately, and later merged, with the software burrow wheeler algorithm (BWA v0.7.15) (Li and Durbin 2009) using the mem algorithm and default parameters to a previously published reference transcriptome of *Bombina bombina* from NCBI (Genbank Accession: HADQ00000000.1). The resultant mapping files were then parsed using Samtools v1.3.1 (Li et al. 2009).

### Mitogenomes

We mapped the processed reads using BWA twice independently to two published reference *B. bombina* mitogenomes, one from Austria (Genbank Accession: JX893173.1) (Pabijan et al. 2013) and the other from Germany (Genbank Accession: MH893761.1) (De Cahsan et al. 2019). Due to insufficient and low read coverage, we removed one sample from the German Högsdorf population (Ge-Ho-KQ01). Mitochondrial consensus sequences were built using a maximum effective base depth approach in ANGSD (Korneliussen et al. 2014) and specifying the following parameters -minMapQ 25 -minQ 25 -uniqueOnly 1. We then compared the total number of mapped reads per individual for each reference (Supplementary Table S1). In order to rule out ascertainment biases, which could be caused by mapping all individuals to a single reference from an ingroup population, we additionally constructed two independent maximum likelihood phylogenetic trees, one for each reference (Supplementary Material, Fig. S2). This was done by computing an alignment of our 29 *B. bombina* samples and two outgroup mitogenome sequences (*B. orientalis* (Genbank accession: AY957562.1), *B. variegata* (Genbank accession: AY971143.1)) for each reference sequence using mafft (Katoh and Standley 2013) and running RAxML (Stamatakis 2006), with 1000 bootstrap iterations, specifying a GTR+G substitution model. The resultant trees were then visually inspected for incongruencies. As the resultant trees were the same regardless of reference, we took the consensus sequence made using the reference that had most reads mapping to it. These were then aligned using mafft. We then produced an unrooted maximum likelihood phylogenetic tree from these aligned consensus sequences by running RAxML with 1000 bootstrap iterations, specifying a GTR+G substitution model.

### Population structure analyses

We performed three independent Principal Component Analyses with the software ANGSD v0.915 and *PCangsd* v0.95 (Meisner and Albrechtsen 2018). We performed two analyses based on identity by state (IBS) and direct SNP calling using ANGSD and the third analysis was based off of genotype likelihoods using ANGSD and PCangsd. We performed one IBS analysis using a single random base per site and one using the consensus base call method. For both IBS analyses we used a minimum base quality (−minQ) and a minimum mapping quality (−minMapQ) parameter of 25, we required 20 individuals as the minimum number for a site to be considered for analysis (−minInd 20), only included reads mapping uniquely to one location (−uniqueonly), removed singletons by only including SNPs found in at least two individuals (−minFreq 0.074), called genotype likelihoods using the GATK method (−GL 2) and only included contigs over 1000 bp in size. We converted the output covariance matrices in PCA inputs using R, which were then plotted using R. For the genotype likelihood method, we produced the PCangsd input file using ANGSD specifying a minimum base quality and a minimum mapping quality parameter of 25, we required 20 individuals as the minimum number for a site to be considered for analysis, only included reads mapping uniquely to one location (−uniqueonly), a minimum minor allele frequency of 0.05 (−minMaf 0.05), called genotype likelihoods using the GATK method (−GL 2), only included contigs over 1000 bp in size and only considered SNPs with a p-value lower than 1e-6 (−SNP_pval). The output genotype likelihoods were then converted into a covariance matrix using PCangsd. The same genotype likelihoods were also used to perform an admixture analysis in PCangsd (−admix). The most probable K value was chosen by the software based on the eigenvalues of the PCA.

### D-statistics or ABBA BABA analysis

We ran D-statistics in ANGSD to look for the degree of admixture in our samples. D-statistics works based on a predefined species tree which uses three ingroup and one outgroup taxon. This topology can be written as (H1,H2),H3),O) where H1 and H2 are more closely related to one another than either are to H3. D-statistics then scans across the genome to find regions that break the known species tree, either due to incomplete lineage sorting or admixture between H3 and either H1 or H2. The occurrence of (H1,H3),H2),O) being equal to (H3,H2),H1),O), is interpreted as only incomplete lineage sorting (or equal amounts of gene flow between H1+H3 and H2+H3). Sites, at which H2 and H3 share a derived allele are called ABBA sites, while BABA represents sites at which H1 and H3 share the derived state. Any deviations from the expected 1:1 ratio is considered as differential levels of gene flow between the ingroup species analysed and is represented by the D value (null expectation of the test: D=0). This value quantifies the deviation from the expected ratio, which is calculated by the differences in the sum of ABBA and BABA site patterns across the genome, divided by their sum:

> *D = [sum(ABBA) – sum(BABA)] / [sum(ABBA) + sum(BABA)]*

We used a random base sampling approach (−do Abbababa 1), considered only transcripts greater than 1000 bp in length, specified *B. orientalis* as outgroup and used the following filters: -minMapQ 25 -minQ 25 -uniqueOnly 1. To investigate the significance of our result, we performed a weighted block jackknife test. D values more than three standard errors different from zero (−3< Z >3) were considered as statistically significant.

## Results

### Mitochondrial genome reconstructions

We obtained 30 complete mitochondrial genomes of *Bombina bombina* from tadpoles sampled in 2016 (Austria: Vienna) and 2017 (Germany: Fehmarn, Eutin/Röbel, Dannau, Högsdorf, Neu-Testorf) using Illumina paired-end 150 bp shotgun sequencing data from cDNA libraries. We performed two independent mapping runs for all samples, using either a German (Genbank Accession: MH893761.1) or an Austrian (Genbank Accession: JX893173.1) mitogenome as a reference. A summary table of the mapping results can be found in Supplementary Table S1. All Austrian individuals show a higher number of mapped reads when mapped to the Austrian reference genome. Four individuals from the German Högsdorf population, as well as one individual from Dannau (Ge-Da-KQ07) also contained higher read numbers mapping to the Austrian reference. All of the other German individuals from Dannau, Eutin, Fehmarn and Testorf contained more reads mapping to the German mitogenome reference than to the Austrian one. To rule out any ascertainment biases that could be introduced by the mapping reference, we initially computed two maximum likelihood trees using RAxML (Stamatakis 2006; Stamatakis 2014), specifying *Bombina orientalis* as an outgroup. One with all individuals mapped to the German reference sequence, and one with all individuals mapped to the Austrian reference sequence (Supplementary Fig. S2). Both trees were congruent in their topology confirming the absence of a bias based on the chosen reference. The bootstrap values support the same two main clades in both trees. We then computed an unrooted tree with RAxML and 1000 bootstrap iterations for all 29 samples using the consensus sequence produced by the reference which gave us the best mapping results for each individual. The resulting tree (Fig. 1) consists of two clades, one including all Austrian individuals, all four German Högsdorf individuals and one German Dannau individual (Ge-Da-KQ07). The other clade includes all German individuals from the populations Fehmarn, Eutin, Testorf and the remaining four individuals from Dannau. The distance between both clades shows a clear separation into a German and an Austrian haplogroup.

**Figure 1:**
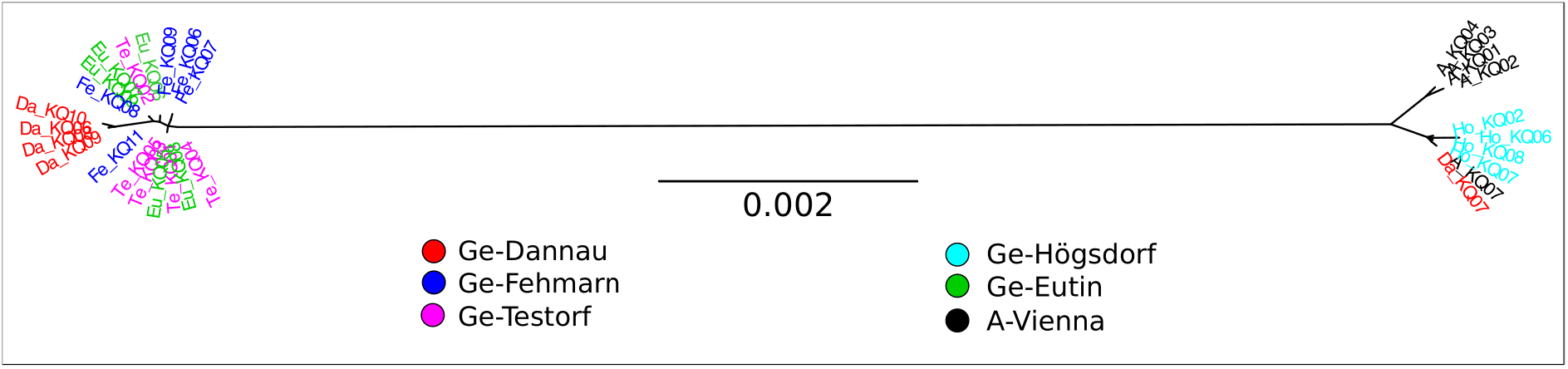
Unrooted Maximum likelihood tree for 29 mitochondrial genomes built with RAxML

### Population structure

To examine whether the pattern found in the mitochondrial genomes are also reflected in the coding regions of the nuclear genome, we investigated 27 transcriptomes for population structure. The same cDNA libraries that we sequenced and used to reconstruct mitochondrial trees, were also used to build transcriptomes for each individual. Mapping was performed with BWA (Li and Durbin 2009) using a published transcriptome of *Bombina bombina* (Genbank Accession: HADQ00000000.1) as a reference. Due to insufficient sequencing coverage, three individuals were removed from the analysis (Ge-Ho-KQ01, Ge-Ho-KQ02, Ge-Te-KQ01), leaving a minimum of three biological replicates per sampling location. We ran a transcriptome wide Principal Component Analysis (PCA) with the software ANGSD to identify population structure. For the analysis we used both identity by state (IBS) and genotype likelihood (GL) methods (see methods). All PCAs produced similar results (Fig. 2 and Supplementary Material, Fig. S3) grouping all Austrian individuals into a single cluster and German individuals from Testorf, Eutin, Dannau and Fehmarn into a second cluster, in agreement with their geographical origin. Interestingly all individuals from Högsdorf form a separate, third cluster. One individual from the German Dannau sampling location (Ge-Da-KQ07) is more closely related to the remaining German individuals but appears to represent an outlier position. This individual carries an Austrian mitochondrial haplotype (cf. Fig. 1).

**Figure 2:**
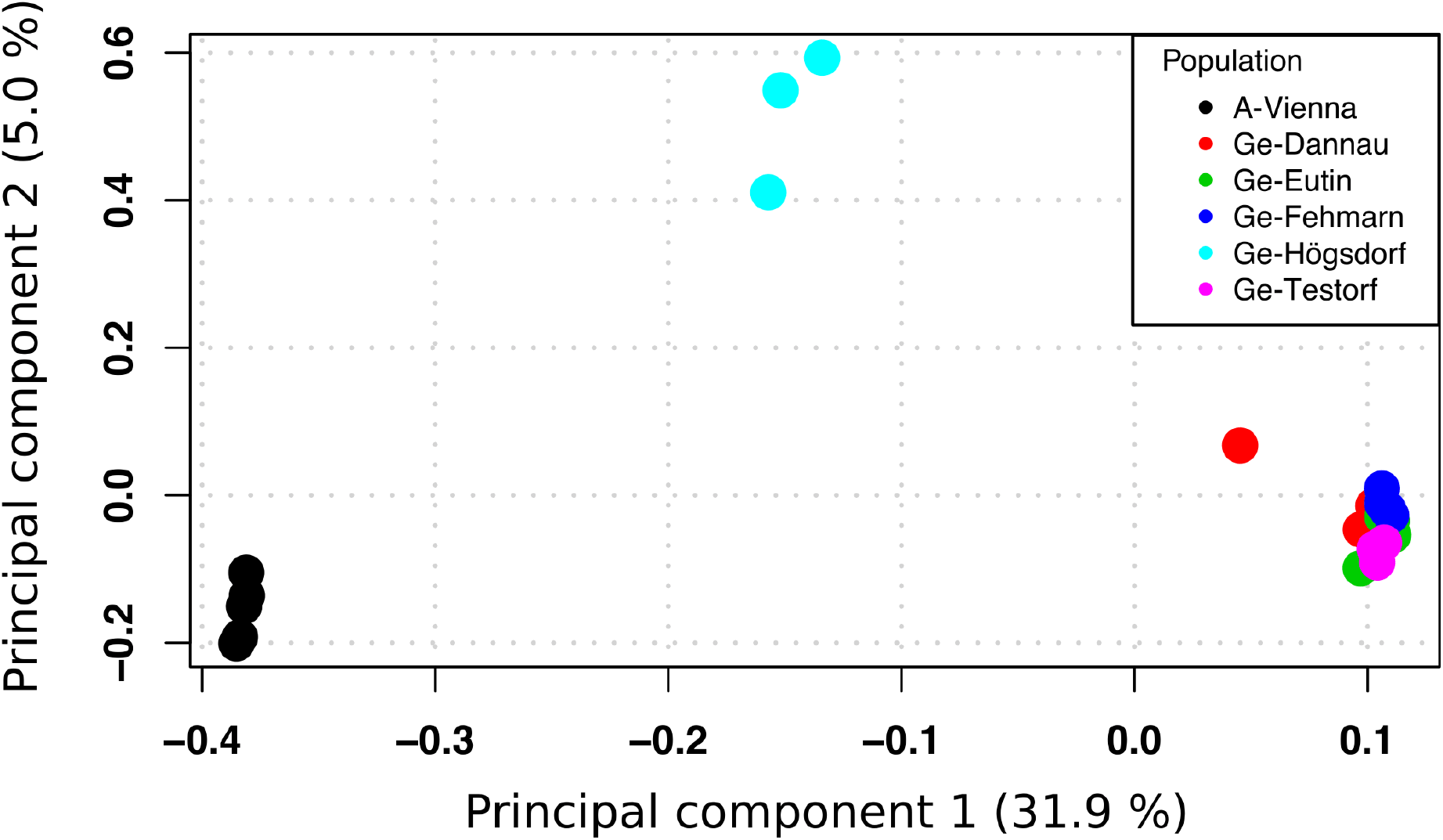
Transcriptome-wide Principal Component Analysis of 27 *B. bombina* tadpole specimen from five locations in northern Germany and one location in Vienna, Austria using the genotype likelihood method (GL)

### Admixture Analysis

In combination, our findings based on the mitochondrial genomes and the PCA on nuclear data suggest some level of admixture between Austrian and German populations. To identify the degree of admixture between southern European fire-bellied toads and northern toads, we performed an admixture structure analysis using PCangsd (Meisner and Albrechtsen 2018). The programme inferred the most likely K value based on the eigenvalues produced during the PCA analysis to be two. The admixture plot shows a high level of shared alleles between all German Högsdorf individuals with Austria, i.e., more than half of the genetic information coming from transcriptome wide SNPs are shared with the Austrian individuals. We also found small levels of Austrian introgression in all German sampling locations except for the island of Fehmarn, where all sampled individuals appear to be uniform German. The second highest level of shared alleles between Austrian and German toads was found in individuals from Dannau, most notably in the tadpole Ge-Da-KQ07. These results are congruent with the results from the PCAs and the mitochondrial haplotype data.

To further evaluate the relative level of introgression between the Austrian and German populations more precisely, we carried out an ABBA BABA or D-statistics analysis for all 27 samples. This analysis compares derived alleles across the transcriptome and investigates these sites for two discordant genealogies from the known species tree. A deviation from an expected 1:1 ratio (ABBA = BABA) for these genealogies could indicate gene flow between the populations of interest or be the result of ancestral population structure. However, ancestral population structure is not the case in our study because northern *B. bombina* populations most likely originated from a single panmictic population that had previously diverged from the Southern lineage (Hofman et al. 2007). For the analysis, we specified *Bombina orientalis* as outgroup and ran it with the software ANGSD. All resulting comparisons with an insignificant Z score (a value between −3 and 3) were removed from further analyses. As previous analyses showed Fehmarn to have no Austrian alleles, we plotted all significant D values for the remaining four German sampling locations while constraining Fehmarn to the H1 branch and Austria to the H3 (Fig. 4A). A positive D value indicates admixture between the H2 and the H3 population, while a negative D value indicates admixture between the H1 and the H3 population.

For all four German populations we found positive D values indicating higher levels of admixture between the Austrian population and the German populations from Testorf, Högsdorf, Eutin and Dannau compared to Fehmarn, ranging from 0.048 to 0.288. The German Högsdorf population has the highest D values from 0.157 to 0.288, followed by Dannau with 0.144-0.194. The Testorf and Eutin populations show similar positive, low D values (around 0.05), indicating a smaller degree of admixture with Austrian toads in these populations. As Austrian fire-bellied toads are known to hybridize with the yellow-bellied toads wherever their distributions overlap (Szymura and Barton 1986; Szymura and Barton 1991; MacCallum et al. 1998), we examined whether signs of admixture with the yellow-bellied toad could be found in our Austrian individuals. For this analysis we placed the Austrian individuals as branch H2, Fehmarn as the H1 population, and a previously published *Bombina variegata* transcriptome as H3. The resulting D values confirm significant admixture between Austrian *B. bombina* and *B. variegata* populations (Fig. 4B). We then investigated whether this admixture between Austrian *B. bombina* and *B. variegata* populations may have crossed into German populations via admixture with Austria. For this analysis we placed the most introgressed German population of Högsdorf as branch H2, Fehmarn was specified as the H1 population, and *B. variegata* as H3. Interestingly, this analysis also shows positive D values, indicating admixture between the German Högsdorf population and the yellow-bellied toad. No significant D scores were found for any other German populations. However, as the ranges of Northern German *B. bombina* populations do not overlap with those of *B. variegata*, this most likely reflects an indirect nuclear introgression of *B. variegata* via Austrian *B. bombina* individuals into Germany (Högsdorf) rather than direct admixture.

## Discussion

Through the recovery of complete mitochondrial genomes from five northern *B. bombina* populations and one Austrian *B. bombina* population, we confirmed the presence of southern “alien” mtDNA (i.e., Austrian) in northern Schleswig-Holstein populations of *B. bombina* as found in previous studies using toads collected in 2006 (Schröder et al. 2012). In addition to this general finding based on mtDNA, we also revealed the extent of this introgression at a much finer scale through the inclusion of transcriptome-wide nuclear data. These nuclear data provide many more loci resulting in a much greater sensitivity to find signals of introgression compared to only mtDNA which is purely maternally inherited as a single locus. This increase in loci enabled us to uncover signs of introgression in a number of northern populations, where admixture from southern *B. bombina* populations was not previously detected. In contrast to previous studies, all of our sampling sites, except for Fehmarn, showed traces of introgression from the South. The finding of the Fehmarn population being the least introgressed was not unexpected as it is located on an island separated from the mainland by the brackish water of the Baltic Sea, preventing the dispersal of toads to or from the island. The only connection with the continent is the Fehmarn Sound Bridge, a one kilometer long, man-made car and rail crossing with intensive traffic representing another physical barrier for amphibians. However, as the Fehmarn population recently recovered from a bottleneck resulting in only 13 individuals in 2004 (Drews et al. 2011), the loss of introgressed alleles via genetic drift, exacerbated by low diversity cannot be ruled out as an alternative, albeit less likely explanation for the lack of introgressed individuals.

Furthermore, we found Högsdorf to contain the highest amounts of introgression from Austria, which is in agreement with a previous study (Schröder et al. 2012). However, due to the lack of resolution, this previous study was unable to confidently explain their finding. This lack of resolution arose as Högsdorf was a newly established population from 2001/2002, where only putatively autochthonous toads from a nearby military training area of Eutin were released. At that time and with only a few genetic markers, no signs of mitochondrial (control region) or nuclear introgression (microsatellites, MHC) were found in samples from Eutin (Schröder et al. 2012). However, using transcriptome-wide data we were able to confirm the presence of admixed loci in Eutin as well. The higher level of introgression found for the three individuals from Högsdorf suggests a significantly greater impact by allochthonous individuals from the Southern lineage in this population. This is also reflected in the presence of the Austrian mitochondrial haplotype in all three Högsdorf samples, confirming previous results that this alien haplotype appears to be fixed in this population.

Although the three individuals from Högsdorf show a large extent of Austrian introgression (Fig. 4A), we were still able to detect a substantial affinity to German populations (Fig. 3) not detected in previous studies. As this population stands out as the most introgressed population in the area, it seems reasonable to conclude that at Högsdorf, Southern toads must have been released in greater numbers, at a higher frequency, or more recently than at any other sampling location.

**Figure 3:**
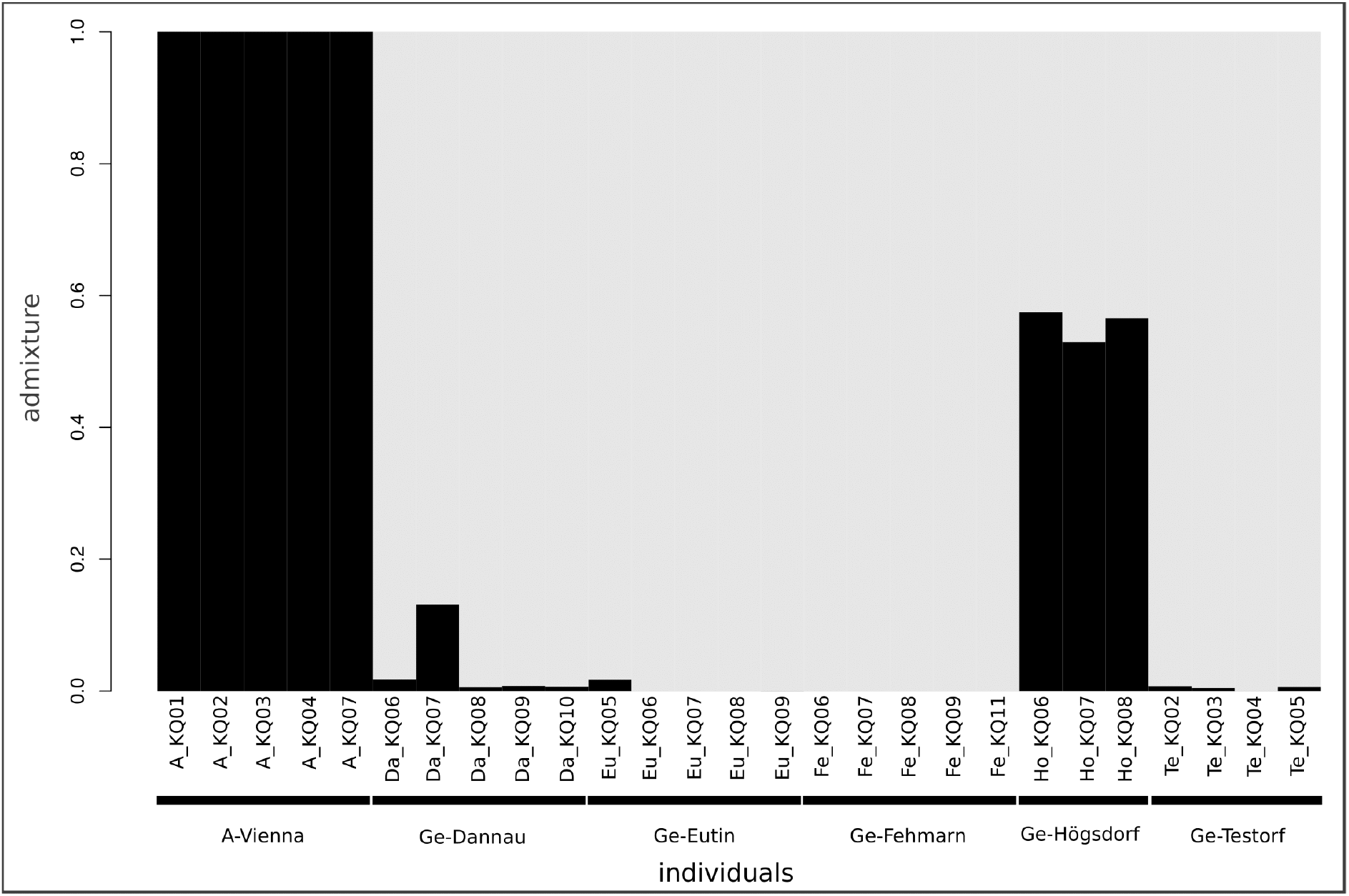
Structure Plot for 27 transcriptomes of German and Austrian *B. bombina* tadpoles (k=2) from six different sampling locations

**Figure 4:**
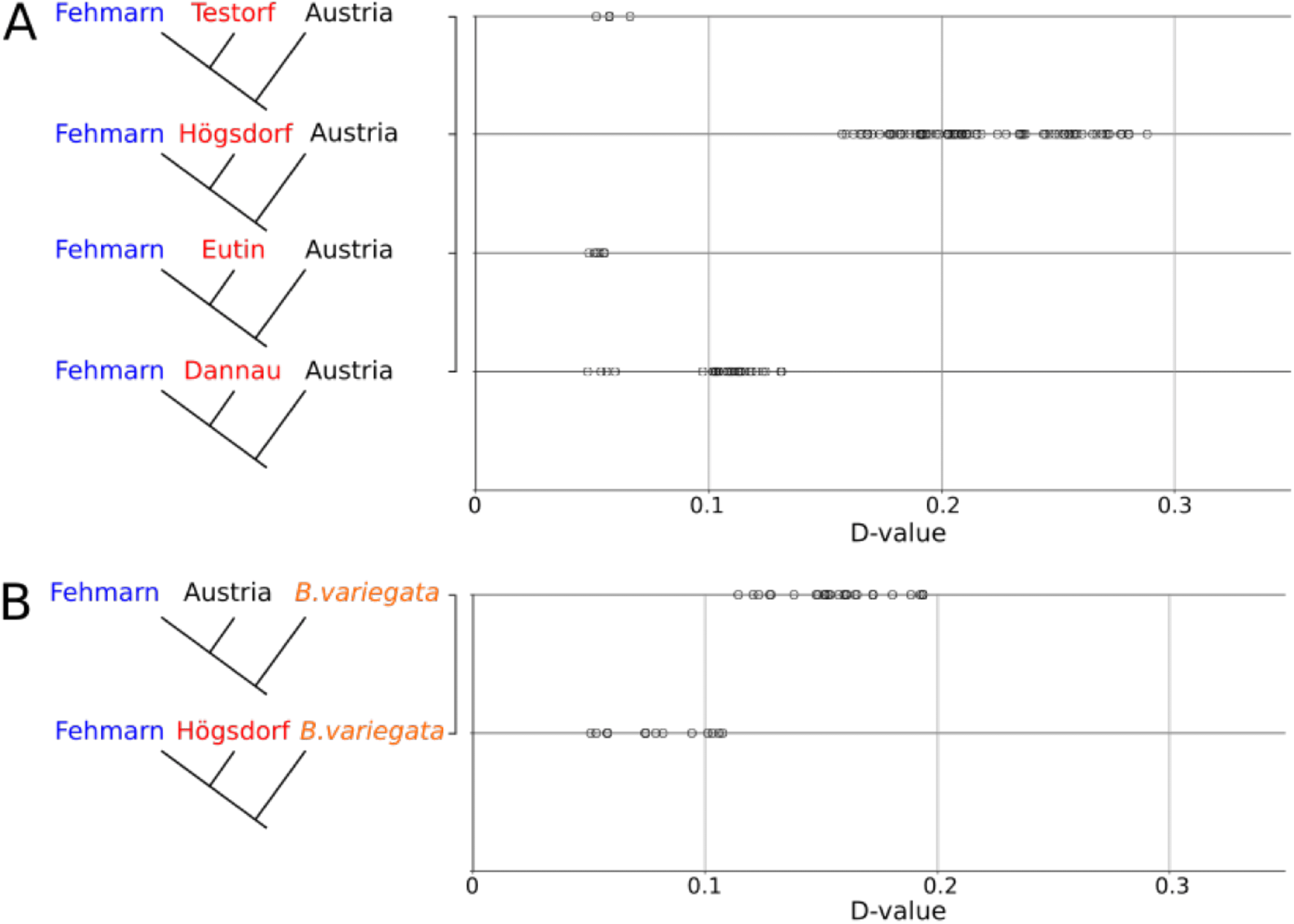
ABBA BABA or D-statistics analysis for 27 transcriptomes of German and Austrian *B. bombina* tadpoles, Outgroup in all analyses is *B. orientalis* (**A**) D values for H1=Fehmarn, H2=Testorf/Högsdorf/Eutin/Dannau, H3=Austria (**B**) D values for H1=Fehmarn, H2=Austria/Högsdorf, *H3=B.variegata*

Additionally, our analysis revealed a significant but less pronounced nuclear introgression for all individuals from Dannau, Eutin and Testorf, despite the southern mitochondrial haplotype only being found in one Dannau specimen (Ge-Da-KQ07). This asymmetric introgression of mitochondrial and nuclear DNA might be explained by genetic drift and the nonrecombinant maternal inheritance of mtDNA (Ballard and Whitlock 2004; Bachtrog et al. 2006; Currat et al. 2008; Zink and Barrowclough 2008). The discrepancy in mitochondrial and nuclear data may also be due to the fact that migrating individuals carrying Austrian genes were mainly males, thus not passing on the Austrian mitochondrial haplotype into adjacent populations. On the other hand, at Högsdorf, where the Austrian mitochondrial haplotype is fixed, Austrian females might have had a higher reproductive success over the German females and/or putatively released Austrian individuals approximately 20 years ago were mainly females.

In contrast to the previous study (Schröder et al. 2012) only finding Högsdorf, out of the populations we studied, to contain introgressed individuals, we find signs of Austrian introgression in almost all of our German populations. This could have a number of explanations. One simple explanation for this could be the increase in number and sensitivity of the chosen marker or pure stochastic chance during sampling. Another scenario is migration of *B. bombina* individuals between introgressed and non-introgressed sites in the last ten years may have spread the introgressed alleles to neighbouring populations. Surprisingly few studies have been published on dispersal in *B. bombina*, despite the prominence of this species in hybrid zone research and its endangered status in its northern distribution area. Adults regularly migrate up to 500 m, and occasionally more than 1 km (Engel 1996; Günther and Schneeweiss 1996). Probably most of the dispersal is accomplished by subadults, which may move over longer distances, like in the better studied sister species *B. variegata* (Gollmann and Gollmann 2012). The greatest distance reported of a natural colonization of nearby ponds for *B. bombina* was in South Sweden (Scania). The colonization rate from the original breeding ponds increased from one to approximately one hundred ponds per year between the years 1983 and 2007, covering distances of 5.5 - 9 km (Drews et al. 2011). The German sampling locations Dannau, Eutin/Röbel and Neu-Testorf are between 5 - 14 km away from the previously uncovered introgressed population Högsdorf, which could explain the newly found signs of nuclear introgression in the adjacent populations. Moreover, conservation efforts, such as the reconstruction of small water bodies, trenches and copses in recent years, may have created sufficient corridors for toad migration throughout the German sampling area. A means behind this migration could have been hearing auditory cues from calling conspecific individuals nearby leading *B. bombina* individuals to gradually migrate to more distant ponds. They could also be drawn to the calls of other anurans, like the tree frog (*Hyla arborea)*, indicating a source of water nearby. Such a process has previously been shown in the great crested newt and the palmate newt (Madden and Jehle 2017).

Finally, as introgressive hybridization between the yellow- and the fire-bellied toad was found in Austria (Gollmann 1984), we tested for admixture between the yellow-bellied toad and the Austrian fire-bellied toad. We confirmed this ancestral gene flow between Austrian *B. bombina* and *B. variegata*. Furthermore, we found *B. variegata* alleles in the highly introgressed Högsdorf population but in none of the other northern German *B. bombina*. We interpret this result as the nuclear genetic information being displaced from *B. variegata* via Austrian *B. bombina* individuals, which were translocated to Högsdorf. This result provides the first evidence of traces of the genetic composition of *B. variegata* being detected in the genepool of an East Holstein *B. bombina* population in northern Germany, outside of the species natural narrow hybrid zone. This result shows that these translocations not only introduced conspecific alleles but also relict alleles of a sister species, the consequences of which are unknown.

The introduction of allochthonous individuals from a genetically distant population led to an increase in genetic variation by introducing novel alleles in four German *B. bombina* populations. Populations at the margin of a species range are often highly fragmented, less diverse in their genetic composition and lack gene flow from surrounding populations (Arnaud-Haond et al. 2006; Bridle and Vines 2007).

Translocated specimens can be a risk to recipient populations, as they may carry foreign parasites/pathogens (Cunningham 1996; Kock et al. 2010) and may disrupt locally adapted gene assemblages by an influx of foreign alleles (outbreeding depression; Edmands 2007). However, an introgression may also be adaptive, as it increases genetic diversity by providing new potentially adaptive gene variants, reducing the level of inbreeding (Edmands 2007; Charlesworth and Willis 2009). We could, show that the translocation of individuals from Austria had a significant genetic impact on this region, with a tendency of spreading further. A positive population trend has been reported for introgressed populations. Whether this is a purely demographic phenomenon or is associated with adaptive introgressed alleles at particular loci, has to be evaluated. Our transcriptomes of autochthonous specimens from two separate evolutionary lineages (Northern and Southern) form a basis to search for expressed lineage-specific SNPs at distinct loci; the fate of the respective alleles could then be assessed transcriptome-wide in the introgressed populations to identify potential adaptive variants. This will unravel loci at which introgressed alleles may be adaptive and may have been favored by selection.

## Acknowledgements

This work was supported by the German Research Foundation (DFG, TI 349-13-1). The authors like to acknowledge Arne Drews at the Landesamt für Landwirtschaft, Umwelt und ländliche Räume for his support regarding sampling permits in Schleswig-Holstein, Germany. We also like to thank Prof. Dr. Michael Lenhard for the access to his computing server, and the High Performance Computing Cluster Orson2, managed by ZIM (Zentrum für Informationstechnologie und Medienmanagement) at the University of Potsdam where large-scale computations were performed.

## Data Accessibility

Newly reported sequences will be uploaded and stated here with accession numbers upon acceptance.

## Author Contributions

The project was conceived by R.T. B.D, M.V.W, H.D, M.O, and G.G assisted with locating and sampling of *B. bombina* tadpoles in the field. The majority of lab work was performed by B.D, including RNA extraction, mRNA enrichment and cDNA library construction, with additional help during library build from S.P; Sequencing was outsourced and performed by the company Novogene (Hong Kong, China); B.D and M.V.W carried out DNA analyses and interpretation of results. Final editing and manuscript preparation was coordinated by B.D. All contributing authors read and agreed to the final manuscript.

