## Supplementary Information for "Introgression of Austrian fire-bellied toads (*Bombina bombina*) into northern German populations confirmed by complete mitochondrial genomes and transcriptome-wide Single Nucleotide Polymorphisms (SNPs)"

### Supplementary Material

**Table S1:** Comparison of total number of mapped reads for each tadpole sample, one *B. variegata* and one *B. orientalis* specimen to either a German (Genbank Accession: MH893761.1) or an Austrian reference mitogenome (Genbank Accession: JX893173.1), three samples marked with an asterisk were excluded from downstream analysis due to low read coverage

| | | | | Mitogenome mapping results | | | Transcriptome mapping results | |
|---|---|---|---|---|---|---|---|---|
| Country of origin | Specimen ID | No. of raw sequencing reads | Percent of combined reads | No. of reads mapped to German reference mitogenome | | No. of reads mapped to Austrian reference mitogenome | No. of mapped reads | Read depth |
| Austria | A_KQ01 | 47159871 | 73.67% | 34958 | < | 35076 | 34460457 | 119 |
| | A_KQ02 | 42720234 | 75.60% | 32701 | < | 32833 | 32326557 | 105.1 |
| | A_KQ03 | 44560704 | 69.83% | 30704 | < | 30790 | 33401245 | 127 |
| | A_KQ04 | 47185408 | 73.33% | 32165 | < | 32248 | 34663772 | 121.4 |
| | A_KQ07 | 48103134 | 77.26% | 35590 | < | 35695 | 35076193 | 112.8 |
| Germany | Da_KQ06 | 49237963 | 71.82% | 32268 | > | 32231 | 25123535 | 80.4 |
| | Da_KQ07 | 48654408 | 68.89% | 35875 | < | 36093 | 29729217 | 92.3 |
| | Da_KQ08 | 53364709 | 72.29% | 32286 | > | 32238 | 25537466 | 84.5 |
| | Da_KQ09 | 44378398 | 68.33% | 29988 | > | 29896 | 24145673 | 91.9 |
| | Da_KQ10 | 45557881 | 74.28% | 32343 | > | 32203 | 25112580 | 89.4 |
| | Eu_KQ05 | 52498185 | 83.05% | 34281 | > | 34167 | 26258941 | 84.7 |
| | Eu_KQ06 | 39215861 | 80.02% | 32511 | > | 32364 | 20492306 | 75.5 |
| | Eu_KQ07 | 53181140 | 73.79% | 28597 | > | 28517 | 23643927 | 79.3 |
| | Eu_KQ08 | 49330157 | 71.80% | 32578 | > | 32445 | 25316598 | 78.5 |
| | Eu_KQ09 | 52823169 | 68.64% | 29124 | > | 29000 | 25358014 | 81.3 |
| | Fe_KQ06 | 59005617 | 73.50% | 31956 | > | 31888 | 31168228 | 106.1 |
| | Fe_KQ07 | 52048554 | 75.92% | 33893 | > | 33786 | 25942907 | 83.5 |
| | Fe_KQ08 | 86140518 | 82.26% | 34634 | > | 34546 | 42132866 | 121.5 |
| | Fe_KQ09 | 50079255 | 69.39% | 30672 | > | 30587 | 25753250 | 88.3 |
| | Fe_KQ11 | 46915955 | 74.13% | 27533 | > | 27499 | 22920629 | 89.1 |
| | Ho_KQ01* | 54703 | 76.62% | 1548 | < | 1550 | 25462 | 3.2 |
| | Ho_KQ02* | 5052227 | 72.63% | 18212 | < | 18243 | 2496362 | 15.8 |
| | Ho_KQ06 | 45691165 | 75.77% | 29041 | < | 29145 | 21132444 | 77.3 |
| | Ho_KQ07 | 41269513 | 76.84% | 26265 | < | 26336 | 18438636 | 75.1 |
| | Ho_KQ08 | 54233242 | 78.78% | 30854 | < | 31022 | 26063766 | 93.2 |
| | Te_KQ01* | 1709037 | 61.33% | 16768 | > | 16750 | 31855008 | 98.3 |
| | Te_KQ02 | 61921295 | 81.03% | 38521 | > | 38333 | 22433162 | 79 |
| | Te_KQ03 | 41276360 | 81.44% | 33530 | > | 33391 | 21405314 | 70.6 |
| | Te_KQ04 | 49154994 | 80.64% | 32699 | > | 32573 | 24068777 | 76.1 |
| | Te_KQ05 | 49298569 | 80.85% | 36741 | > | 36617 | 38067686 | 75 |
| Poland | *B.variegata* (AY971143.1) | 55301570 | 89.32% | 41960 | > | 41900 | 45720738 | 95.9 |
| Korea | *B.orientalis* (AY957562.1) | 46932624 | 81.73% | 24530 | > | 23819 | 990210 | 11.4 |

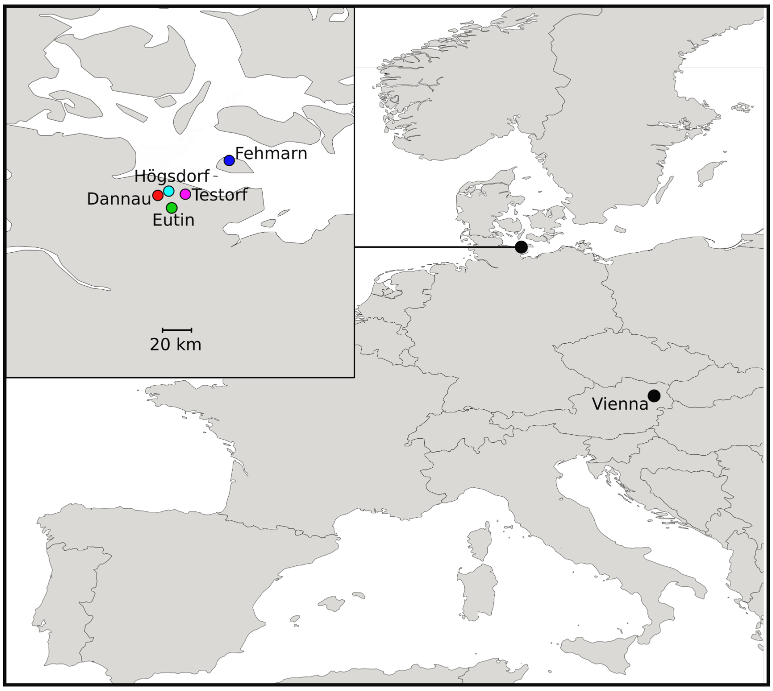


**Figure S1:** Map of sampling locations in Germany (5) and Austria (1)

**A**

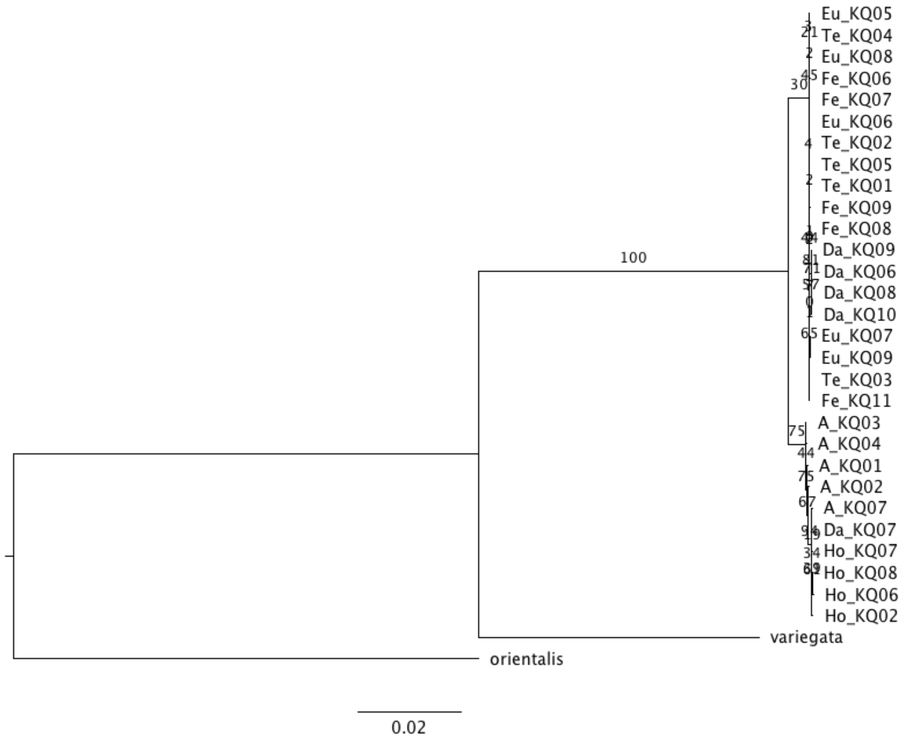


**B**

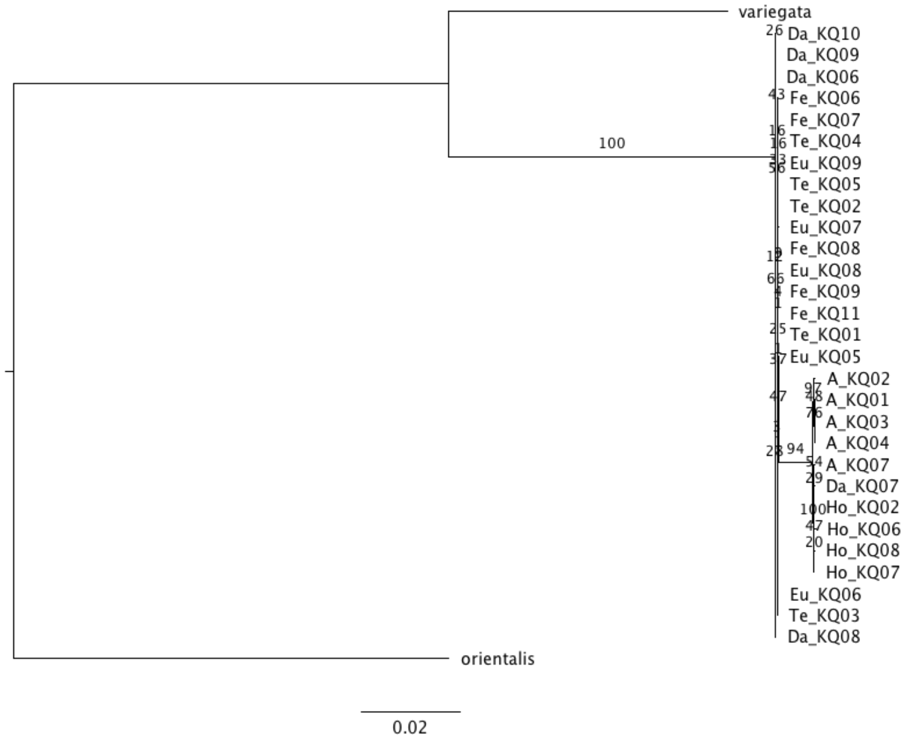


**Figure S2:** Maximum Likelihood trees of 29 mitogenomes mapped to (A) a German reference mitogenome or (B) an Austrian reference mitogenome using RaxML specifying *B. orientalis* as outgroup. Numbers on branch lengths represent bootstrap values.

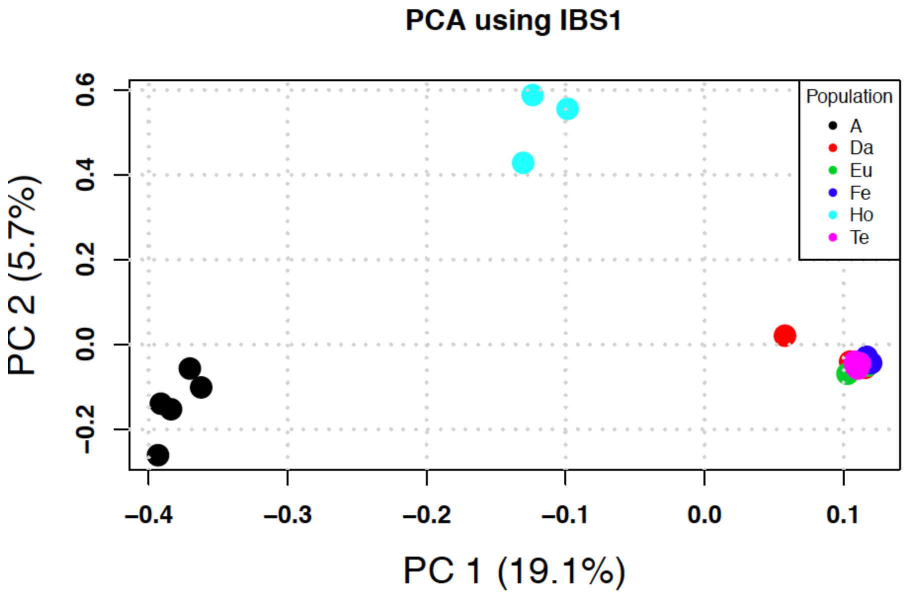


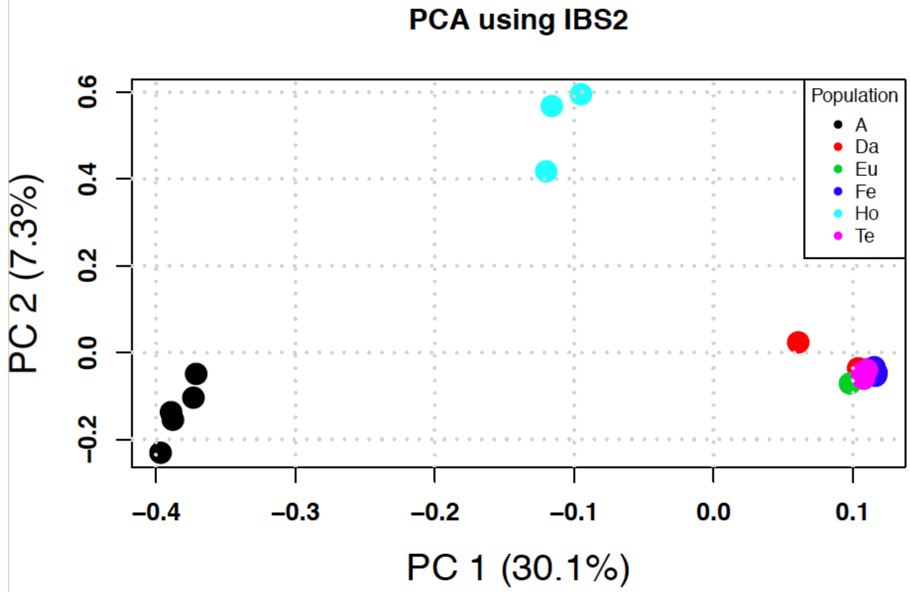


**Figure S3:** Transcriptome-wide Principal Component Analysis of 27 *Bombina bombina* tadpole specimen from five locations in northern Germany and one location in Vienna, Austria using the identity by state (IBS) method
